# Core genes can have higher recombination rates than accessory genes within global microbial populations

**DOI:** 10.1101/2021.09.13.460184

**Authors:** Asher Preska Steinberg, Mingzhi Lin, Edo Kussell

## Abstract

Recombination is essential to microbial evolution^1–4^, and is involved in the spread of antibiotic resistance, antigenic variation, and adaptation to the host niche^5–8^. Yet quantifying the impact of homologous recombination on different gene classes, which is critical to understanding how selection acts on variation to shape species diversity and genome structure, remains challenging. This is largely due to the dynamic nature of bacterial genomes, whose high intraspecies genome content diversity^9^ and complex phylogenetic relationships^10–12^ present difficulties for inferring rates of recombination, particularly for rare genes. In this work, we apply a computationally efficient, non-phylogenetic approach^13,14^ to measure homologous recombination rates in the core and accessory genome (genes present in all strains and only a subset of strains, respectively) using >100,000 whole genome sequences from 12 microbial species. Our analysis suggests that even well-resolved sequence clusters sampled from global populations interact with overlapping gene pools, which has implications for the role of population structure in genome evolution. We show that in a majority of species, core genes have shorter coalescence times and higher recombination rates than accessory genes, and that gene frequency is often positively correlated with increased recombination. Our results provide a new line of population genomic evidence supporting the hypothesis that core genes are under strong, purifying selection^15–17^, and indicate that homologous recombination may play a key role in increasing the efficiency of selection in those parts of the genome most conserved within each species.

## Main

Bacterial genomes contain both a set of genes with key functions that are found in nearly all strains of a given species, known as the ‘core’ genome (defined here as genes found in >95% of strains), and a collection of genes found in only a subset of strains referred to as the ‘accessory’ genome^9^. Inherent to bacterial genome evolution is the process of homologous recombination, in which fragments of DNA taken up from the environment are incorporated into homologous sites in the genome, often obscuring clonal signals and confounding phylogenetic analyses^10,11,18^. While it is known that bacterial genomes recombine to such an extent that it is unclear if their evolution can be characterized as tree-like^10,19,20^, the ability to quantify recombination rates in different parts of the genome has been limited. Particularly, for accessory genes, frequent gene loss and gain events^21–23^, in addition to sparse data on rare genes, cause difficulties in applying standard phylogeny-based approaches to infer recombination rates^24–31^. Measuring homologous recombination rates in different parts of the genome will contribute to our fundamental understanding of how the interplay between selection and recombination shapes sequence diversity, which has important implications for how bacteria adapt to stresses such as antibiotics, host immune responses, and nutrient conditions. Here, we determine how homologous recombination rates differ between core and accessory genes.

To infer the parameters of homologous recombination, we apply and extend the *mcorr* method^13,14^, an approach that avoids phylogenetic reconstruction by using a coalescent-based population genetics model with recombination to capture the statistics of large-scale sequencing data (see ref.^13^ for details). In the simplest form of the model, we define a ‘sample’ to be a set of lineages with mean coalescence time *T*_*sample*_, which mutate and exchange homologous DNA fragments with a much larger ‘pool’ of sequences having mean coalescence time *T*_*pool*_ (Fig. 1A). Different samples can access distinct pools (Fig. 1A) or, alternatively, these pools can have significant overlap leading to DNA exchange between each sample and multiple pools (Fig. 1B). By fitting the model to the data, we infer recombination parameters of both the sample and its pool. Moreover, by examining a range of samples from a given species, we can understand how recombination has affected different samples and the diversity of their pools. In this work, we focus primarily on three parameters: the pool’s mutational divergence, *θ*_*pool*_ ≡ 2*μT*_*pool*_, the pool’s recombinational divergence, *ϕ*_*pool*_ ≡ 2*γT*_*pool*_, and the sample’s recombination coverage, *c*_*sample*_ (where *μ* and *γ* are the synonymous substitution rate and recombination rate, respectively). Together, *θ*_*pool*_ and *ϕ*_*pool*_ estimate the number of synonymous substitutions and recombination events that have occurred per site since coalescence of the pool, while *c*_*sample*_ estimates the fraction of the genome that has been replaced by homologous recombination with the pool since coalescence of the given sample.

**Fig 1.**
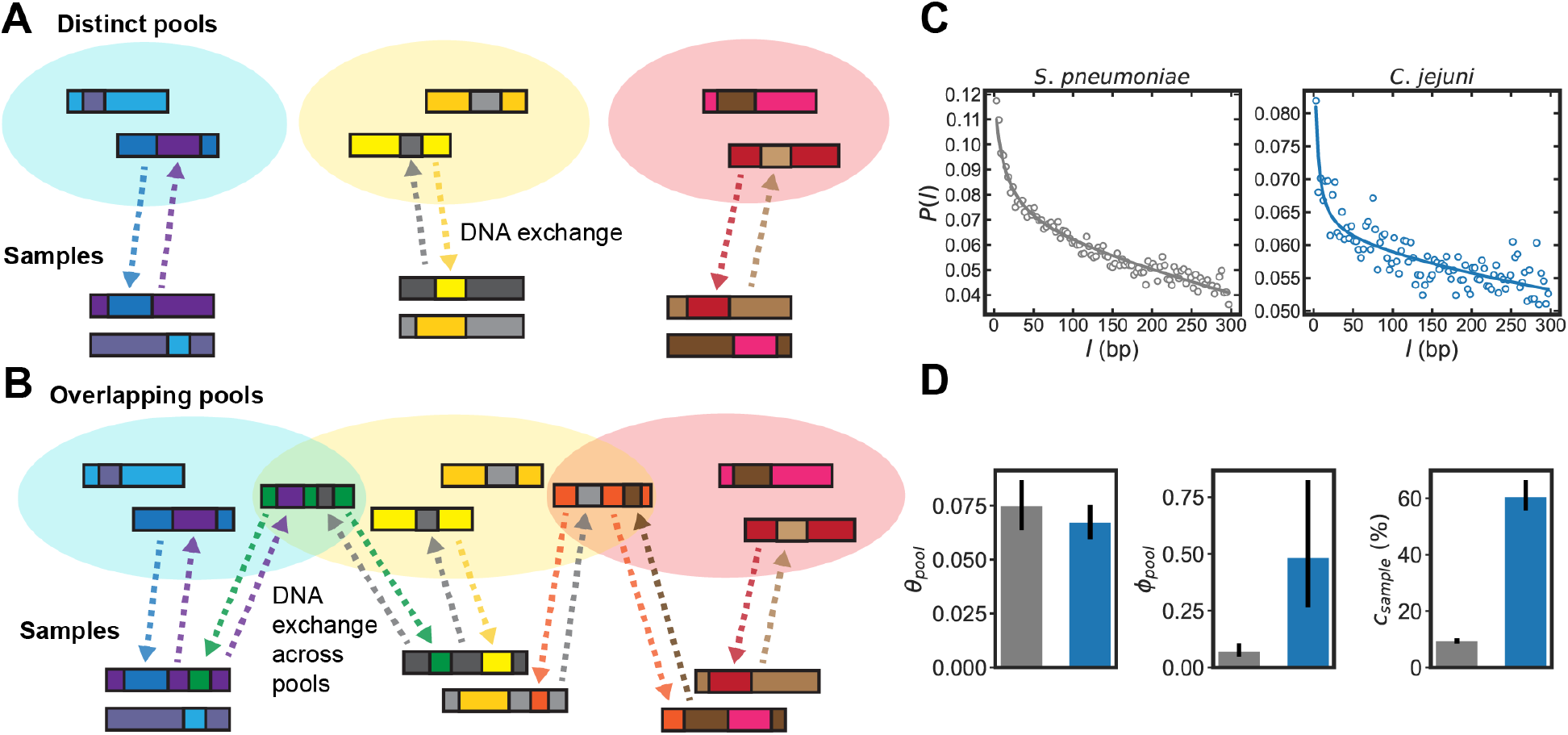
Inferring parameters of homologous recombination from whole genome sequences. (A,B) Schematics depicting exchange of homologous DNA fragments between sets of lineages (“samples”) and larger “pools” of DNA sequences. In panel A, each sample interacts with a distinct pool, while in panel B, samples exchange DNA with multiple, overlapping pools. (C) Example correlation profiles of synonymous substitutions for the core genes in samples consisting of 568 *Streptococcus pneumoniae (S. pneumoniae)* and 2215 *Campylobacter jejuni (C. jejuni*) WGS (further description of these datasets are given in the following sections). (D) Recombination parameters inferred from fitting the profiles shown in panel C to a population genetics model (see Methods and ref. ^13^ for details). Left to right: the pool’s mutational divergence (*θ*_*pool*_), the pool’s recombinational divergence (*ϕ*_*pool*_), and the sample’s recombination coverage (*c*_*sample*_). Error bars are 95% bootstrap confidence intervals (see Methods for details). Colors correspond to panel C.

The model predicts the conditional probability of a difference at genome site *i+l* given a difference at site *i*, which we refer to as the ‘correlation profile’, *P(l)*, where *l* is the distance between sites in basepairs (bp). We take a set of whole genome sequences (WGS) to constitute a sample, and use alignments of protein coding (CDS) regions to measure profiles of synonymous substitutions for all possible sequence pairs, yielding an average profile for the sample. For each pair of sequences in the sample at genome position *i*, a binary variable *σ*_*i*_ is assigned 1 for a difference or 0 for identity. The correlation profile is given by *P*(*l*) ≡ *P*(*σ*_*i*+*l*_ = 1|*σ*_*i*_ = 1), where *i* is restricted to third-position sites of codons. Fitting the model to these correlation profiles yields the parameters of homologous recombination described above (see ref. ^13^ and Methods for details of model and fitting); flat profiles indicate a lack of recombination, while in the presence of recombination *P*(*l*) has a monotonic decay with *l*, and declines more rapidly with increasing recombination^14^. Example profiles and inferred parameters are shown in Fig. 1C-D using sets of *Streptococcus pneumoniae* (*S. pneumoniae*) and *Campylobacter jejuni* (*C. jejuni*) WGS; these sequences are part of the larger datasets used in the analysis that follows. Thus, the method infers parameters of both the sample and the pool of sequences with which it has recombined without using or inferring any phylogenetic relationships, offering a key advantage in determining recombination rates across accessory genes.

To study recombination across the core and accessory genome of a given bacterial species, we sought to understand the recombination behavior of a discrete set of samples from the global microbial population. Samples may have access to different sequence pools, e.g., due to geographic constraints or differences in host niches. The distribution of pool parameters inferred from different samples indicates how different the pools are in terms of their diversity (as schematized in Fig. 1A-B). To delineate a discrete set of samples for a given species, we cluster WGS solely based on sequence similarity using the average linkage algorithm^32^ with the pairwise synonymous diversity (*d*_*s*_) as the distance metric; such approaches are widely used to define well-resolved genotypic clusters within bacterial populations without the inference of phylogenetic relationships^33^. We then analyze each cluster as a separate sample, as well as pairs of clusters, to infer recombination parameters both within and between their pools. To quantify parameters within each pool, we analyze individual clusters and fit correlation profiles measured by averaging over all of a given cluster’s sequence pairs, which yields the parameters of the pool that the cluster interacts with. Specifically, we obtain the *within-pool* divergences, *θ*_*pool*_(*w*/*n*) and *ϕ*_*pool*_(*w*/*n*), as well as the amount of recombination that has taken place within a cluster since coalescence (*c*_*sample*_). To infer parameters between pools, we take pairs of clusters and fit correlation profiles measured by averaging over all sequence pairs consisting of one sequence from each of the two clusters. This yields the *between-pool* divergences, *θ*_*pool*_(*btwn*) and *ϕ*_*pool*_(*btwn*), which estimate the number of substitution and recombination events per site, respectively, since coalescence of both pools with each other. We then infer the degree of overlap between pools using paired comparisons of between- and within-pool divergences. Moreover, by analyzing all possible pairs of clusters, we examine diverse collections of lineages as our samples, yielding a parameter distribution that accounts for the full range of potential interactions between lineages, regardless of their distance in sequence space and agnostic to population structure or sampling biases.

We began by analyzing *S. pneumoniae*, and used the *PubMLST* database (see Supplementary Table 4) which includes WGS of strains from across the globe. For each WGS, we aligned all sequencing reads to a reference genome to create an alignment of all CDS regions (see Methods for details). We then measured *d*_*s*_ across all genes (i.e., core and accessory) for each strain pair, clustered based on these distances to make a dendrogram, and split the sequences into flat clusters, where no sequence within a cluster was more distant than the 10^th^ percentile of pairwise distances, which corresponded to *d*_*s*_ = 0.015 (Fig. 2A, Methods for details; mean *d*_*s*_ values within and between clusters for core and accessory genes are given in Supplementary Table 2). This resulted in 44 major clusters (where a major cluster has >100 strains) encompassing 24,097 strains.

**Fig. 2.**
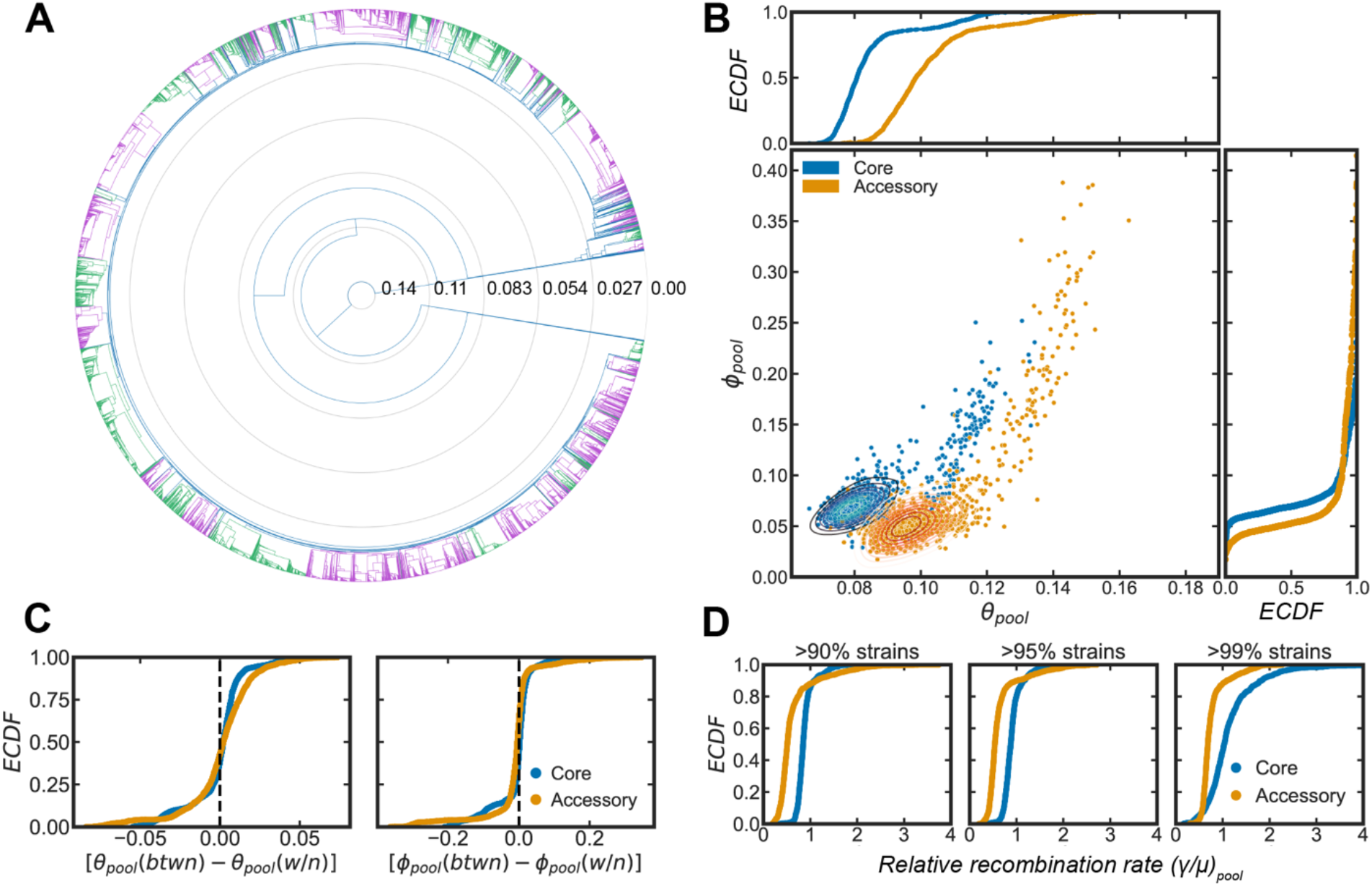
Inference of recombination parameters for the core and accessory genome across a set of *S. pneumoniae* sequence clusters. (A) Dendrogram resulting from hierarchical clustering of 26,599 whole genome sequences (WGS) from the *PubMLST* genome collection for *S. pneumoniae* using the average linkage algorithm. The dendrogram was cut at the 10th percentile of all measured pairwise distances (*d*_*s*_ ~ 0.015) yielding a discrete set of flat clusters. Alternating colors delineate adjacent clusters in the dendrogram. (B) Recombinational divergence (*ϕ*_*pool*_) and mutational divergence (*θ*_*pool*_) inferred from correlation profiles measured using the 44 major clusters of *S. pneumoniae* strains for the core and accessory genome. Each point corresponds to a single within-pool or between-pool measurement; in the main plot, data are overlaid with kernel distribution estimates (smoothed with a Gaussian kernel) to indicate density of observations. Marginal plots show empirical cumulative distribution functions (*ECDFs*) of *θ*_*pool*_ and *ϕ*_*pool*_. Core genes are defined as genes found in >95% of strains. We used model selection with the Akaike Information Criterion to ensure that each profile was well-fit (see Methods for details). (C) *ECDF*s of the paired differences of *θ*_*pool*_ for between-pool (*btwn*) and within-pool (*w/n*) divergences. For each point, [*θ*_*pool*_(*btwn*) − *θ*_*pool*_(*w*/*n*)] = *θ*_*pool*_(*i*, *j*) − *θ*_*pool*_(*i*), where *θ*_*pool*_(*i*, *j*) is the between-pool divergence inferred using clusters *i* and *j*, and *θ*_*pool*_(*i*) is the within-pool divergence inferred using cluster *i*. If either *θ*_*pool*_(*i*) or *θ*_*pool*_(*j*) could not be inferred, the paired differences for [*θ*_*pool*_(*i*, *j*) − *θ*_*pool*_(*i*)] and [*θ*_*pool*_(*i*, *j*) − *θ*_*pool*_(*j*)] were excluded. The same analysis was performed for *ϕ*_*pool*_. (D) *ECDFs* of the relative recombination rates, (*γ*/*μ*)_*pool*_, for the core and accessory genome for all between- and within-pool measurements using different indicated thresholds to define core genes.

We inferred the within- and between-pool parameters for the core and accessory genomes of *S. pneumoniae* using all major clusters (946 total cluster pairs) and plotted these parameters against each other (Fig. 2B). We found that the core and accessory genes of *S. pneumoniae* have distinct distributions of *θ*_*pool*_ and *ϕ*_*pool*_ (Fig. 2B), indicating these gene classes have different evolutionary dynamics. We examined the overlap of sequence pools by plotting the paired differences of the between- and within-pool parameters (Fig. 2C). If the set of pools is distinct, the between-pool divergences should be larger than within-pool divergences. Instead, we find that for *S. pneumoniae*, the paired differences of between- and within-pool divergences are tightly distributed around zero, suggesting that the population structure of *S. pneumoniae* consists of a set of strongly overlapping gene pools (as schematized in Fig. 1B). An alternative hypothesis is that this perceived overlap is the result of where the dendrogram was cut; namely that more finely resolved sequence clusters would exhibit distinct pools. However, when we cut the dendrogram at the 1^st^ and 0.5^th^ percentiles of pairwise distances (*d*_*s*_ = 0.0045, 0.0018, respectively), we observe that the paired differences are generally distributed about zero, again indicating overlapping pools (Supplementary Fig. 1).

As an additional control, we tested how the parameters changed when more accessory genes were included by building a reference pangenome from multiple *S. pneumoniae* genome assemblies using *Roary*^34^, and aligning to this and measuring correlation profiles (Supplementary Fig. 2; Methods). The observed trends were similar, and while this alignment method included more genes (6200 versus 2018), we found that it greatly reduced the aligned percentage per gene (64% versus 98%). We therefore chose to align to a single reference genome for subsequent analyses.

Overall, we observed two main differences between core and accessory genes. First, *θ*_*pool*_ is lower in core versus accessory genes. This indicates that core genes have shorter mean coalescence times than accessory genes, a conclusion that holds as long as the synonymous substitution rate *μ* does not differ systematically between core and accessory genes. While *μ* could potentially vary across the genome due to effects such as codon usage bias^35^, long term microbial evolution experiments have found that *μ* is fairly homogeneous across the core and accessory genomes^36,37^. Second, using the fact that *ϕ*_*pool*_/*θ*_*pool*_ = *γ*/*μ*, the dependence on coalescence time can be removed to yield the relative recombination rate in the sequence pool (from hereon we will append the subscript *pool* to indicate this, i.e., (*γ*/*μ*)_*pool*_). The inferred distribution of (*γ*⁄*μ*)_*pool*_ shows that for *S. pneumoniae*, the core genome has a higher recombination rate than the accessory genome (Fig. 2D). Further, as there are various thresholds used for the percentage of strains which must have a gene for it to be considered “core”^9,34,38,39^, we tested how the parameter distributions shifted with different thresholds and found similar trends (Fig. 2D). We next sought to understand these trends in a broader, multi-species context.

We expanded our analysis to >100,000 genomes from 12 different microbial species (Fig. 3). Using the procedure described in the previous section to cluster sequences and measure correlation profiles within and between clusters, we inferred parameters for the global sequence pool of each species. As with *S. pneumoniae*, we examined the paired differences of between- and within-pool recombination parameters (Supplementary Fig. 3) and found a range of behaviors, which reflect differences in the global pool structures (i.e., overlapping or distinct pools) for each species. Some species (e.g., *S. pneumoniae, H. influenzae*) appeared to have highly overlapping global gene pools, while others had distinct, structured sets of pools (e.g., *H. pylori*). We then analyzed paired differences of core and accessory genome parameter values for all clusters and cluster pairs. Core-accessory pairs were plotted against each other (Fig. 3A-C) and as distributions of paired differences (Fig. 3D-F). Additionally, we performed the Wilcoxon signed-rank test on each paired difference distribution (Supplementary Table 1).

**Fig. 3.**
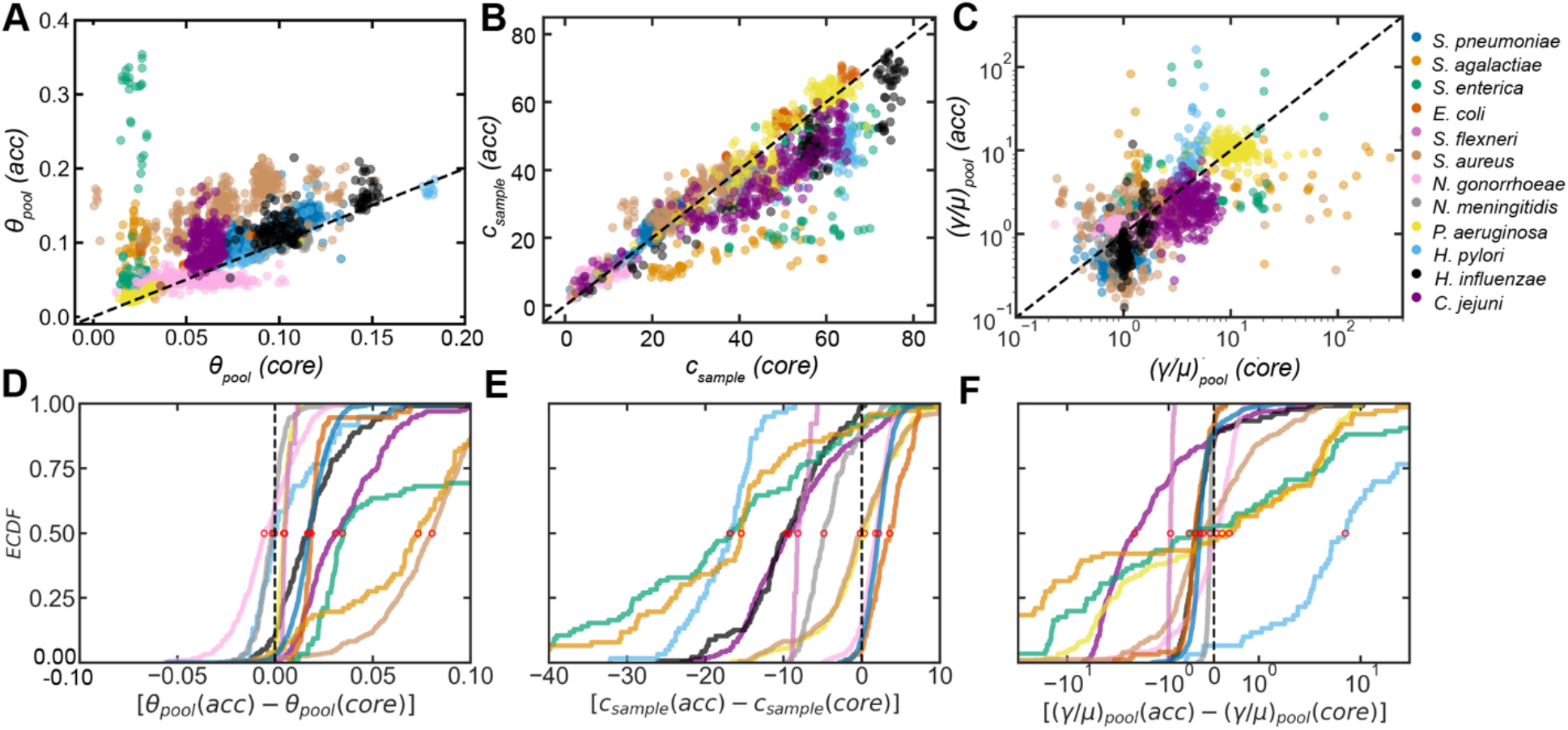
Paired recombination parameter distributions for 12 microbial species. **(A-C)** Scatterplots of inferred parameters for core/accessory genome pairs for individual clusters and pairs of clusters in each microbial species. Values for accessory genomes are plotted on the vertical axis (*acc*) and core (*core*) on the horizontal. Dashed line is y = x. *θ*_*pool*_ and (*γ*/*μ*)_*pool*_ are unitless, *c*_*sample*_ is given as the percentage of recombined genomic sites. **(D-F)** Empirical cumulative distribution functions (*ECDFs*) of paired differences for each core/accessory genome pair for the inferred parameters. Dashed line indicates where the paired difference equals zero. Red open-face markers indicate the median of each distribution. The x-axis of panel F is on a symmetric log-scale. Two-sided p-values were calculated using the Wilcoxon signed-rank test to compare the paired distributions for each parameter (Supplementary Table 1). When model selection was performed with AIC (described in Methods) if part of a core/accessory pair was poorly fit, the paired sample was excluded. Full species names, number of strains, number of clusters, and mean synonymous diversity within and between clusters are reported for each species in Supplementary Table 2. Supplementary Figs. 4–5 show *ECDF*s of recombination parameters for the core and accessory genomes plotted separately (i.e., not as paired differences).

We found that across species *θ*_*pool*_ was generally higher for the accessory genome (Fig. 3A,D), indicating that mean coalescence times are shorter for the core genome. The core genome has been seen to exhibit stronger purifying selection than the accessory genome^9,15–17,40^, and it is well-known that purifying selection reduces mean coalescence times^41,42^. An additional consideration is that some accessory genes are used for niche specialization^9,39,43^; if different alleles of the same accessory gene are adaptive to different niches, this could result in allelic diversity due to balancing selection, which would also increase coalescence times. Therefore, the shorter coalescence times of the core genome are consistent with the hypothesis that core genes are under stronger purifying selection, and additionally may reflect that a subset of accessory genes are under diversifying selection.

In a majority of species, both *c*_*sample*_ (Fig. 3B,E) and (*γ*/*μ*)_*pool*_ (Fig. 3C,F) were significantly higher for the core genes (see Supplementary Table 1 for results of Wilcoxon signed-rank test), indicating higher rates of homologous recombination. While it is well known that accessory genes exhibit both high gene replacement rates and high sequence variability^21,44–46^, gene replacement generally occurs by non-homologous recombination mechanisms, and rates of homologous recombination in the accessory genome have previously been difficult to assess. One hypothesis consistent with our observations is that homologous recombination homogenizes and preserves genes with critical functions found in the core genome^21^. We note that because higher levels of recombination in core genes are expected to reduce the effects of background selection^42^, the relative reduction in *θ*_*pool*_ observed in core versus accessory genes likely underestimates the difference in strength of purifying selection acting on these gene classes. In certain species, mechanisms that give rise to higher recombination rates in the core genome are known; for example, some members of the *Neisseriaceae* (e.g., *N. meningitidis* and *N. gonorrhoeae*) and *Pasteurellaceae* families (e.g., *H. influenzae*) have DNA uptake sequences and uptake signal sequences accumulated in their core genomes which promote homologous recombination^47,48^. A mutually compatible, alternative hypothesis is that the accumulation of mutations in the accessory genome (evidenced by their higher *θ*_*pool*_) leaves these genes with less homology than core genes, reducing the rate of homologous recombination.

One testable hypothesis consistent with our observation that core genes generally undergo more homologous recombination when compared with accessory genes is that the least frequent genes experience less recombination because they have fewer recombination partners. We tested this by binning the genomes for each species into four gene frequency classes consistent with prior delineations of the pangenome as follows (*f* is frequency across strains):^34,49^ “cloud” genes (*f* ≤ 15%), “shell” genes (divided into two segments, 15% < *f* ≤ 55%, 55% < *f* ≤ 95%), and core genes (*f* > 95%). We then inferred the distributions of recombination parameters within each gene class for each of the 12 microbial species (Fig. 4A-B). We found that certain species displayed trends consistent with the gene frequency hypothesis, particularly for the recombination coverage (Fig. 4A). When considering the recombination rates (displayed as empirical cumulative distribution functions here because of disparate rates across and within species), *C. jejuni*, *S. agalactiae*, *P. aeruginosa*, and *E. coli* showed trends most consistent with this hypothesis. For some of the exceptions (e.g., *N. gonorrhoeae*, *N. meningitidis*), the cloud genes showed the highest recombination rates, which could be related to the high gene replacement rate experienced by rare genes such as “ORFans”^44^, or diversifying selection experienced by these genes. Overall, we see a modest positive correlation with both recombination coverage and rates when considering the 50^th^ and 25^th^ percentiles of the matched parameter distributions for all species (Fig. 4C-D).

**Fig. 4.**
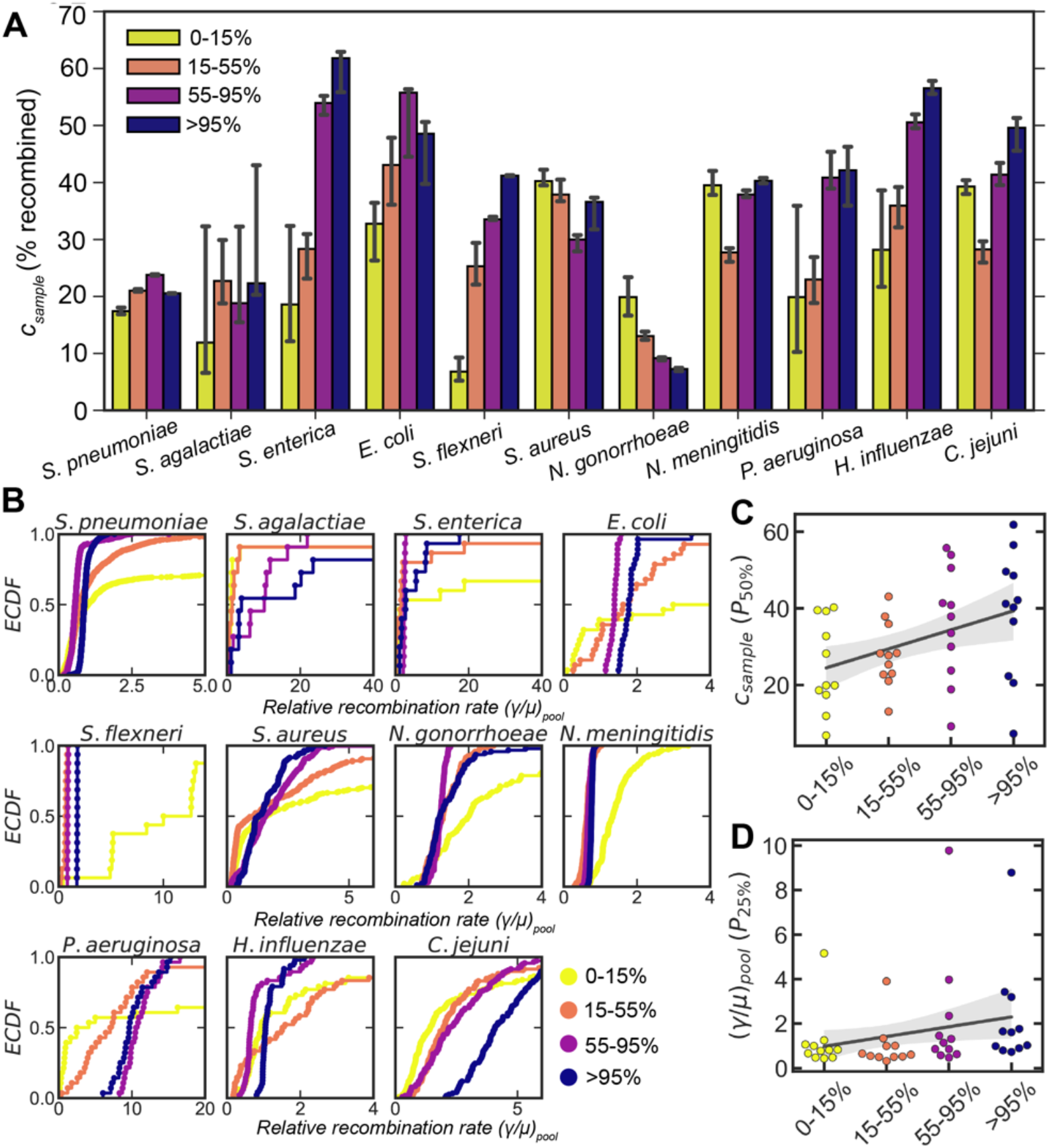
Dependence of recombination rates on gene frequency. (A) Bar plots depicting the medians of the recombination coverage (*c*_*sample*_, given as the percentage of recombined sites) for genes of different frequency (i.e., the percentage of strains which have a given gene) for eleven microbial species. Matched parameter distributions were inferred from correlation profiles within and between clusters for each species. Error bars are 95% bootstrap CIs measured by randomly sampling with replacement from the parameter distributions. 10,000 bootstrap replicates were calculated. Supplementary Fig. 6 shows corresponding empirical cumulative distribution functions (*ECDF*s) for each species. (B) *ECDF*s *of* the relative recombination rate, (*γ*/*μ*)_*pool*_, for different gene frequencies corresponding to the same matched distributions shown in *A*. Two-sided p-values were calculated using the Friedman test to compare the matched distributions shown in *A* and *B* (Supplementary Table 3). (C-D) The 50th percentile of the recombination coverage (*C*) and the 25th percentile of the rate (*D*) distributions for each species plotted against gene frequency. The solid gray line is a simple linear regression; shaded areas are 95% CIs of the regression estimated by bootstrapping the medians for each species. P-values were calculated using the Wald test, where the null hypothesis is that the slope is less than zero; *p* = 0.003 and 0.05 for panels *C* and *D*, respectively. For all panels, only matched samples where parameters could be inferred for each gene frequency bin were included (i.e., all correlation profiles were deemed well-fit by model selection using AIC as described in Methods). For *H. pylori* this excluded almost all matched samples (< 3 remained), leaving us with insufficient data to perform this analysis. For each frequency range, upper bounds are inclusive, lower bounds exclusive.

There is appreciable interest in understanding why microbes have pangenomes and, further, how different parts of the genome have evolved^9,39,44,50^. While it is known that homologous recombination plays a key role in shaping the genome^1,2,4,21,51^, study of this aspect of genome evolution has been limited. This was due to both computational bottlenecks and a reliance on phylogenetic-based methods^24–31^, the latter of which hindered determination of recombination parameters for accessory genes, whose phylogenies are challenging to ascertain. Here, we expanded our non-phylogenetic, computationally-efficient *mcorr* framework to overcome these obstacles, and present a comprehensive analysis using >100,000 genomes from 12 microbial species. Our analysis suggests that while for some species the global gene pool consists of highly structured sets of well-defined pools, for other species the boundaries between pools are ‘blurred’, and the global gene pool appears to be a set of highly overlapping pools. Recent analyses have suggested that some microbial species have highly structured populations in which population structure plays a key role in shaping evolutionary dynamics^12,52,53^; our work complexifies this picture, indicating that some species have highly interconnected evolutionary dynamics regardless of their population structure. This further emphasizes the need for future investigations on the role of population structure in microbial genome evolution.

We found that coalescence times are generally shorter for core genes, offering genomic data in support of the hypothesis that core genes are under strong, purifying selection^9,15–17,40^. Further, we found that despite the high gene replacement rates and sequence variability experienced by some accessory genes^21,44–46^, core genes generally have higher levels of homologous recombination than accessory genes in a majority of species. As we also observed that core genes generally had lower mutational divergences, their increased homology may reduce the barrier for homologous recombination, leading to higher rates of recombination in the core genome. Thus, we hypothesize that an evolutionary feedback loop may operate in certain species of bacteria: purifying selection acting on core genes causes higher levels of homology at these loci, this increases their homologous recombination rates, which in turn enables more efficient purifying selection by breaking linkages between genes, effectively minimizing Hill-Robertson interference and hitchhiking effects^54–56^ across the core genome. Further investigations are needed to examine whether such an evolutionary mechanism can act to fine-tune recombination rate variation across microbial genomes. Moreover, we found evidence which is generally consistent with the hypothesis that the levels of homologous recombination experienced by a given gene are correlated with its prevalence within a given microbial species. Our work yields fundamental knowledge needed to decipher why these broad gene classes (i.e., core and accessory) have evolved, particularly by contributing to our understanding of how selection and recombination shape diversity in these parts of the genome. The expansion of this approach to analyze more specific gene classes (e.g., drug-resistance, metabolism) could lead to a better understanding of the interplay of selective pressure and homologous recombination.

## Methods

### Data and code availability

- Lists of SRA accession numbers corresponding to the raw reads used to build the multi-sequence alignments analyzed in this manuscript will be included as supplemental files to this manuscript. All SRA files, reference genomes, and complete genome assemblies are available through NCBI. All sequence collections used are listed in Supplementary Table 4. For the PubMLST sequence collections, PubMLST was used to identify whole genome sequences (by filtering for strains in the “Genome Collection” of each species where the sequence length is at least that of the reference genome), then the raw reads were downloaded from NCBI using their SRA numbers. Accession numbers for reference genomes used for each microbial species are also listed in Supplementary Table 4.
- All original code has been deposited at GitHub and will be made publicly available as of the date of publication. Links are given below:

- https://github.com/kussell-lab/mcorr
- https://github.com/kussell-lab/mcorr-clustering
- https://github.com/kussell-lab/ReferenceAlignmentGenerator
- https://github.com/kussell-lab/PangenomeAlignmentGenerator
- Any additional information required to reanalyze the data reported in this paper is available from the lead contact upon request.

#### Description of coalescent-based population genetics model with recombination

The details of the model derivation and parameters are given in the “Supplementary Notes” of ref.^13^. Here, we provide the model equations necessary for the fitting procedure used in this work.

The measured synonymous diversity of the sample, *d*_*s*_, is a linear combination of the pool diversity, *d*_*p*_ ≡ *d*(*θ*_*p*_) (due to recombination), and the clonal diversity of the sample, *d*(*θ*_*s*_), as follows:

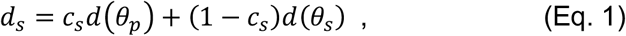

where *c*_*s*_ is the recombination coverage in the sample, and *d*(*θ*) is given by the classic population genetics expression for heterozygosity^57^ (see table below). The predicted form of the correlation profile is

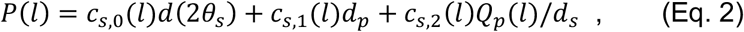

where the functional forms of *c*_*s,i*_(*l*) (*i* = 0,1,2) and *Q*_*p*_ (*l*) are given in the table below. As described in ref. ^13^, we can re-express Eq. 2 in terms of the three parameters *θ*_*s*_, *ϕ*_*s*_, and 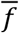, determined by fitting the profiles. From these, the pool parameters are obtained using the following relations 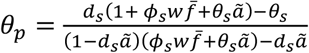 and *ϕ*_*p*_ = *θ*_*p*_*ϕ*_*s*_/*θ*_*s*_; see table below for values of constants 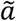 and *w*.

| Term | Expression | Description |
| --- | --- | --- |
| $\theta_s$ or $\theta_p$ | | Mutational divergence of the sample or pool |
| $\phi_s$ or $\phi_p$ | | Recombinational divergence of the sample or pool |
| $\bar{f}$ | | Mean fragment size of homologous recombination |
| $l$ | | Distance (bp) between two loci |
| The following constants are used below: $\tilde{a} = 4/3$ and $w = 2/3$. | | |
| $d(\theta)$ | $\frac{\theta}{1 + \theta \tilde{a}}$ | Synonymous diversity (heterozygosity) for divergence $\theta$ |
| $c_s$ | $\frac{\phi_s w \bar{f}}{1 + \theta_s \tilde{a} + \phi_s w \bar{f}}$ | Recombination coverage of the sample |
| $c_{s,1}(l)$ | $\frac{2\phi_s w l}{1 + 2\theta_s \tilde{a} + \phi_s w (\bar{f} + l)}$ | Probability that the most recent event was a recombination affecting only one locus |
| $c_{s,2}(l)$ | $\frac{\phi_s w(\bar{f} - l)}{1 + 2\theta_s \tilde{a} + \phi_s w(\bar{f} + l)}$ | Probability that the most recent event was a recombination affecting both loci |
| $c_{s,0}(l)$ | $1 - c_{s,1} - c_{s,2}$ | Probability that the most recent event at either locus was not a recombination |
| $Q_p(l)$ | $2 \left( \frac{\theta_p}{1 + \theta_p \tilde{a}} \right)^2 \left( \frac{1 + \theta_p \tilde{a} + \phi_p w l}{1 + 2\theta_p \tilde{a} + 2\phi_p w l} \right)$ | Probability of a difference at both loci in the pool |

#### Calculation of correlation profiles

For a given sample of aligned sequences, we measured the substitution profile, *σ*_*i*_(*k*, *g*), for each pair of sequences *k*, at each position *i* along each gene *g*, where *σ*_*i*_(*k*, *g*) = 1 for a difference and *σ*_*i*_(*k*, *g*) = 0 for identity. In all calculations below, we only consider positions *i* that are third-position sites of codons yielding synonymous substitutions. We computed the pairwise synonymous diversity of each gene *g* as:

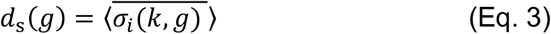

where the bar denotes averaging over all sequence pairs *k* and the bracket denotes averaging over all positions *i*. The joint probability of synonymous substitutions for two sites separated by a distance *l* was calculated for each gene as:

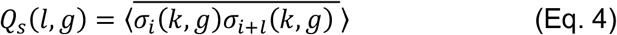

We averaged these over all genes to obtain the *sample diversity*, *d*_*s*_ = (1/*n*_*g*_) ∑_*g*_ *d*_*s*_(*g*), and the joint probability of substitutions at two sites in the sample, *Q*_*s*_(*l*) = (1/*n*_*g*_) ∑_*g*_ *Q*_*s*_(*l*, *g*), where *n*_*g*_ is the total number of genes. The correlation profile is then given by *P*(*l*) = *Q*_*s*_(*l*)/*d*_*s*_.

For calculations of *within-pool* parameters, all possible sequence pairs within a cluster were considered. This was done with the original command-line (CLI) program *mcorr-xmfa* program in the *mcorr* package which takes as input a single extended multi-fasta (XMFA) file. For calculations of *between-pool* parameters using pairs of clusters, only sequence pairs where each sequence was from a different cluster were considered. This was done with the CLI program created for this paper called *mcorr-xmfa-2clades*, which uses two XMFA files (one from each sequence cluster) as inputs. These can be found in the *mcorr* GitHub repository: https://github.com/kussell-lab/mcorr.

#### Fitting of correlation profiles and model selection using Akaike information criterion (AIC)

The basic fitting procedure used was previously described in ref. ^13^. We used the python package LMFIT^58^ version 0.9.7 (https://lmfit.github.io/lmfit-py/) to fit *P*(*l*) to the analytical form given in Eq. 2; in this work, we used the least-squares minimization with Trust Region Reflective method instead of the default Levenberg-Marquardt algorithm and increased the maximum number of function evaluations from 10^4^ to 10^6^ As described in the text, as an additional test of goodness of fit, we compared the fit of the data with Eq. 2 where all parameters vary freely (which we refer to as the “full-recombination model”), to the “null-recombination model” in which we set *c*_*s*,1_ = *c*_*s*,2_ = 0, which yields 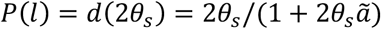, i.e., a constant correlation profile which is fit by taking the average over *l* of the measured values, yielding a single parameter, *θ*_*s*_. We perform model selection between the null-recombination and full-recombination models by evaluating the Akaike information criterion (AIC) for each model, then computing the Akaike weight for each model, which can be roughly interpreted as the probability that a given model yields the best prediction of the data.^59,60^ The AIC was computed using LMFIT using _*AIC*_ = *n* ln(χ^2^/*n*) + 2*N*_*v*_, where *n* is the number of datapoints, *N*_v_ is the number of parameters being varied in the fitting, and 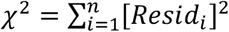 is the sum of the squared residuals for each data point *i*. The Akaike weight for model *j* ∈ {∅, *r*}, can then be calculated as:

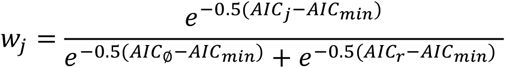

where *AIC*_∅_ and *AIC*_*r*_ denote the AIC values for the null and full-recombination models respectively, and *AIC*_*min*_ = min {*AIC*_∅_, *AIC*_*r*_}. The evidence ratio, which corresponds to the likelihood of one model being favored over the other in terms of minimizing Kullback-Leibler discrepancy^59^, can be computed as *w*_*r*_/*w*_∅_, where *w*_*r*_ and *w*_∅_ are the Akaike weights of the full and null-recombination models respectively. Fits to correlation profiles where *w*_*r*_/*w*_∅_ < 100 were not included in the analyses presented in the main text. We also did not include fits to correlation profiles which yielded unphysical parameter values (*θ*_*pool*_ < 0, *ϕ*_*pool*_ < 0), or extreme values of recombination coverage where parameter fitting becomes less reliable due to too few recombination events (*c*_*sample*_ < 1%) or the over-fitting of a flat profile (*c*_*sample*_ > 99%). The CLI program *mcorrFitCompare* was built for this study to fit correlation profiles to both models, and can be found in the *mcorr* GitHub repository: https://github.com/kussell-lab/mcorr.

#### Clustering using pairwise synonymous diversity

In this study, we extended the *mcorr* method to measure the synonymous diversity separately for each sequence pair, denoted by *d*_*pair*_. This is the same as the calculation of *d*_*s*_ described in the above section “Calculation of correlation profiles” without averaging over sequence pairs *k*. The CLI program *mcorr-pair* measures *d*_*pair*_ for all possible sequence pairs from an XMFA file. We also built a CLI program called *mcorr-pair-sync*, which calculates a subset of all pairwise diversities from a given set of sequences. We primarily relied on the latter program, as it allows for parallelization of jobs on a high performance computing (HPC) cluster. The CLI programs *mcorr-dm* and *mcorr-dm-chunks* were built to collect outputs from *mcorr-pair* and *mcorr-pair-sync* respectively and write them to a square distance matrix for use with standard clustering algorithms. Lastly, we built a program (*clusterSequences*) relying on the python *scipy* package to cluster sequences using the unweighted pair group method with arithmetic mean (UPGMA) or average linkage method with *d*_*pair*_ as the distance metric. All necessary code to perform these calculations can be found in the github repository: https://github.com/kussell-lab/mcorr-clustering.

#### Generation of multi-sequence alignments for core and accessory genes using whole genome sequences

For all collections of whole genome sequences (Supplementary Table 4), with the exception for *H. pylori* (where the multi-sequence alignment was already assembled), we used reference-guided alignment to build consensus genomes from raw reads for each sequence, then extracted the CDS regions of each gene to make extended multi-fasta (XMFA) files. For the collections from the PubMLST database, we identified whole genome sequences by filtering for all sequences in the “Genome Collection” for a given organism which had lengths greater than or equal to the length of the reference genome. We then exported tables which included the corresponding SRA numbers for each strain, and downloaded the reads from NCBI using the corresponding SRA. For the other sequence collections used, we used their associated NCBI Bioproject ID to obtain the SRA numbers for each strain, and downloaded reads in the same way. Reads were mapped against a reference genome (listed in the Supplementary Table 4 for each microbial species) using SMALT (version 0.7.6; https://www.sanger.ac.uk/tool/smalt-0/), and consensus genomes were created using SAMtools mpileup.^61^ We then extracted the alignments of CDS regions from these consensus genomes using genomic coordinates provided from the general feature format (GFF) file of the reference genome to create an XMFA file of CDS regions for each gene. We filtered out any gene alignments where the gene sequence was ≥ 2% gaps. We then measured the percentage of the entire strain collection which had each gene (based on presence/absence of a gene sequence) to determine if a gene should be considered core or accessory. Code to perform this analysis is provided in the GitHub repository: https://github.com/kussell-lab/ReferenceAlignmentGenerator.

#### Generation of multi-sequence alignment to pangenome reference genome for Streptococcus pneumoniae

We downloaded all complete genome assemblies from NCBI for *Streptococcus pneumoniae* (*S. pneumoniae*) as of December 18^th^, 2020 (81 genomes), and used Prokka^62^ to re-annotate the assemblies and generate general feature format version 3 (GFF3) files for use with Roary^34^. Roary was then used to generate a multi-FASTA for each gene CDS region in the pangenome (which we refer to as the “pangenome reference”). We aligned reads to this pangenome reference in the same manner described in “*Generation of multi-sequence alignments for core and accessory genes using whole genome sequences*”, then collected each gene alignment to create an XMFA file for the pangenome. We split the XMFA into files for the core and accessory genome by measuring the percentage of strains which had each gene. In this case, because alignment quality was not as high as when we aligned reads to a single reference genome (as described in the main text), we counted a gene as present if there was only a partially aligned sequence for the gene, and absent if there was no aligned sequence. Code to perform this analysis can be found in the following GitHub repository: https://github.com/kussell-lab/PangenomeAlignmentGenerator.

### Statistical analysis

For the 95% bootstrap CIs appearing in Fig. 1D, the same procedure was used as in ref.^13^. In brief, bootstrap replicates of the set of genes were created by resampling the list of all genes in the genome with replacement, then recalculating *d*_*s*_, *Q*_*s*_, and *P*(*l*) for each replicate. We then performed the same fitting procedure which was done on the actual dataset to infer recombination parameters on each of the bootstrap replicates to generate 95% confidence intervals. The resampling was done 1000 times.

For all other 95% bootstrap CIs (Fig. 4 and Supplementary Figs.), the procedure used for bootstrapping can be found in the Figure legends of the corresponding Figures, and was done using the python package *seaborn*. The process of model selection using AIC is described in the above section “*Fitting of correlation profiles and model selection using Akaike information criterion (AIC)*”. The number of major clusters and sequences used for each microbial species are given in Supplementary Table 2.

## Acknowledgments

This work was supported in part by NIH grant R01-GM-097356. Asher Preska Steinberg is a Simons Foundation Awardee of the Life Sciences Research Foundation. We gratefully acknowledge Nobuto Takeuchi for useful discussions and feedback on the manuscript; the New York University (NYU) high performance computing cluster for resources, and its staff for technical support.

## Author Contributions

A.P.S. and E.K. designed the research, interpreted the results, and wrote the manuscript. M.L. contributed to research design and provided preliminary analyses. A.P.S. performed the bioinformatic analyses, data analysis, wrote additional code for the *mcorr* package and for the processing of sequence alignments used in this paper.

## Supplemental Information

**Supplementary Fig. 1.**
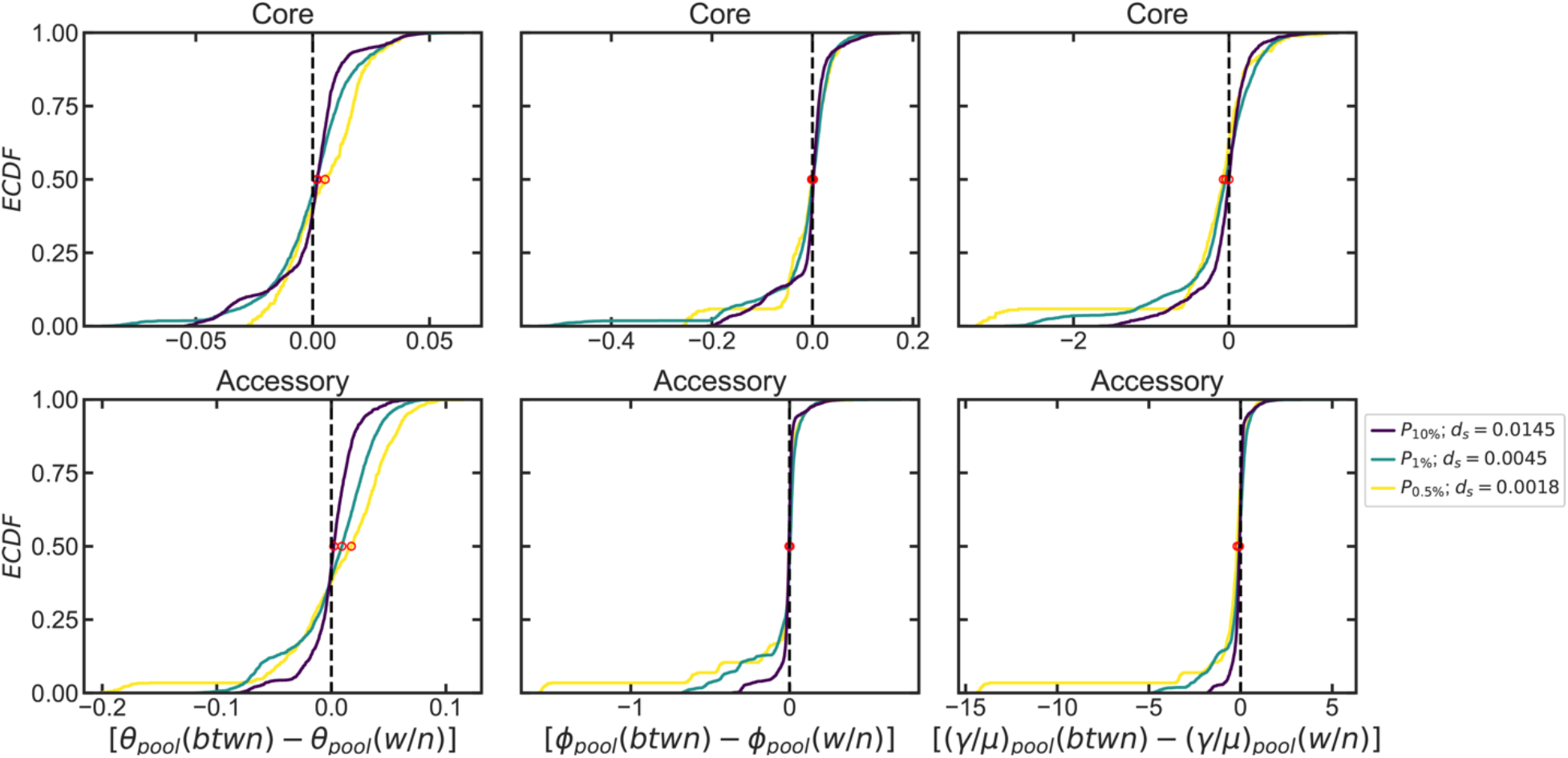
Paired differences of recombination parameters for *Streptococcus pneumoniae* for different minimal distances between sequences in a given cluster. Empirical cumulative distribution functions (*ECDFs*) for paired differences of between-pool (*btwn*) and within-pool (*w/n*) mutational and recombinational divergence (*θ*_*pool*_ and *ϕ*_*pool*_, respectively) and relative recombination rate of the pool ((*γ*/*μ*)_*pool*_, explained in main text). After clustering sequences using the average linkage algorithm, the dendrograms were cut at different percentiles of all pairwise distances (in the legend, *P*_*X*%_ indicates a cut made at the X^th^ percentile of distances, and *d*_*s*_ indicates the corresponding distance value). Only paired differences where the within-pool parameter could be computed for both clusters used to compute the between-pool divergence were plotted. Red circles indicate medians.

**Supplementary Fig. 2.**
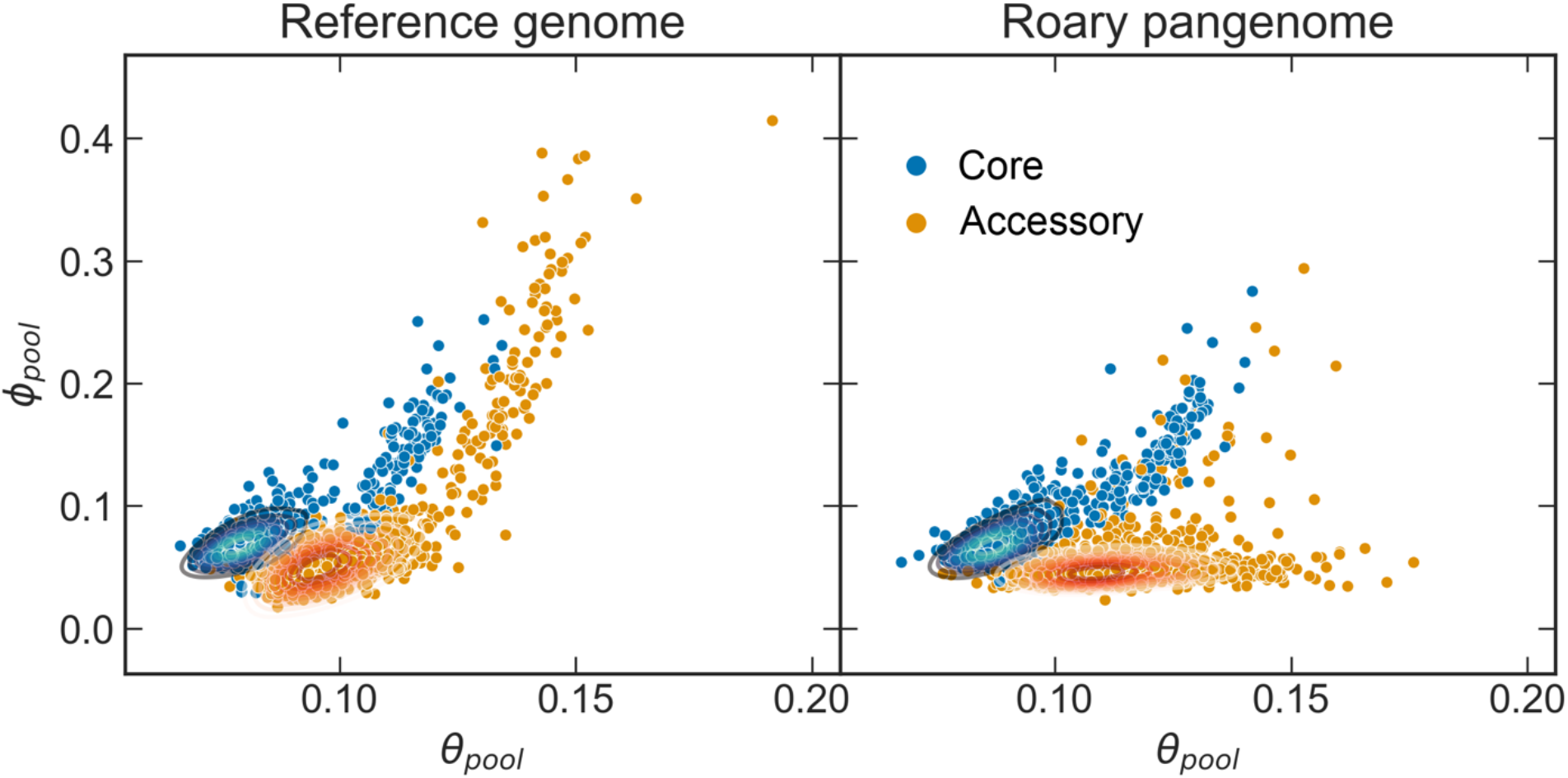
Comparison of distributions of recombination parameters for the core and accessory genome of *S. pneumoniae* with different alignment methods. Recombinational divergence (*ϕ*_*pool*_) plotted against mutational divergence (*θ*_*pool*_) inferred from correlation profiles measured within and between clusters for both the core and accessory genome for each of the major sequence clusters (>100 strains) of ~26,000 *S. pneumoniae* strains from the *PubMLST* collection when raw reads were aligned to a single reference genome (left) compared to when raw reads were aligned to a multi-FASTA of all gene sequences in a pangenome generated by Roary^34^ (right). Each panel shows a scatterplot overlaid with kernel distribution estimates (smoothed with a Gaussian kernel) depicted as contours representing density of observations. Core genes are defined as genes found in >95% of strains. Left plot is reproduced from Fig. 2A. Clustering with the alignment using a single reference genome resulted in 44 major clusters, clustering with the pangenome alignment resulted in 46 major clusters.

**Supplementary Fig. 3.**
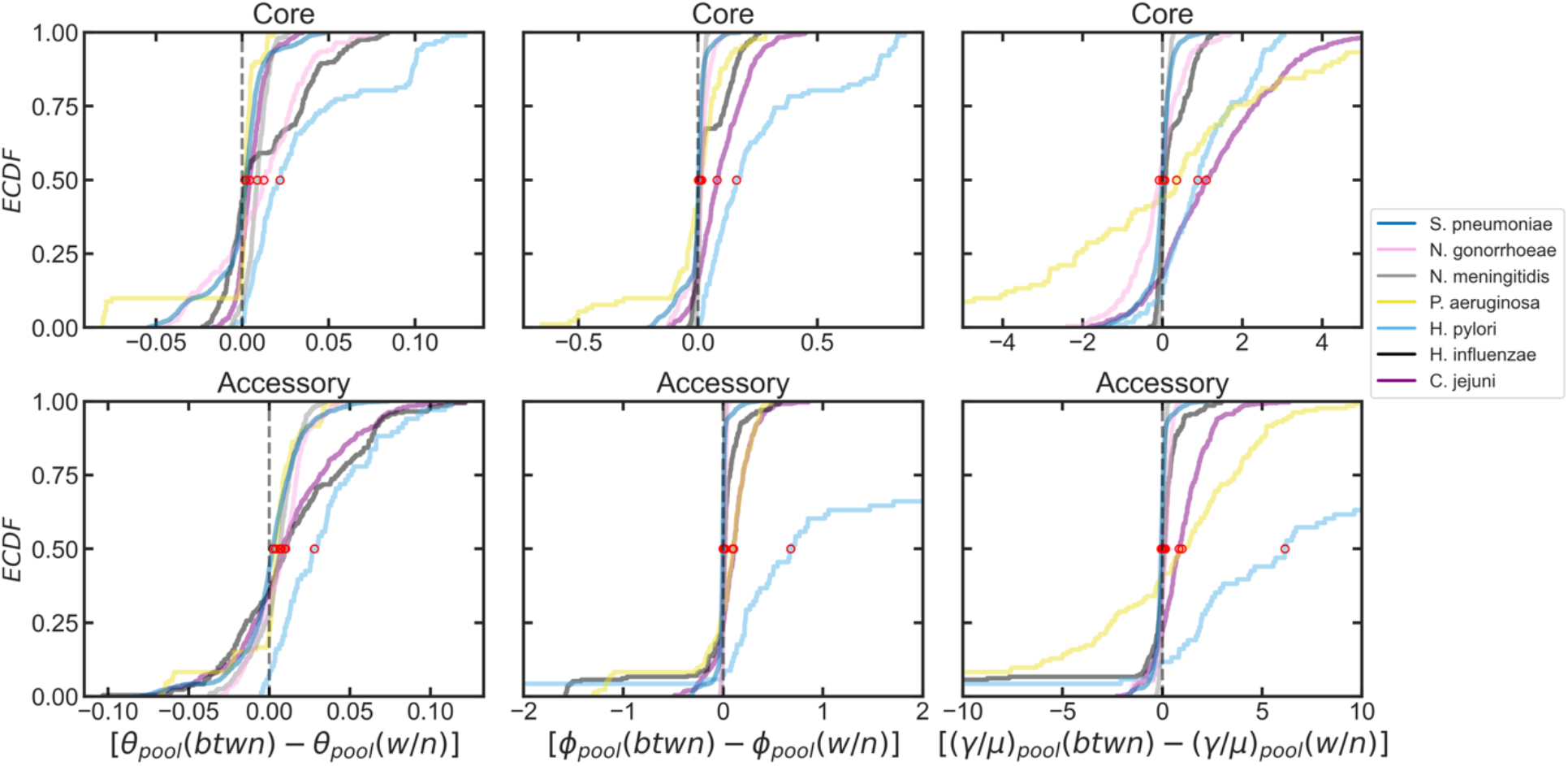
Paired differences of between- and within-pool recombination parameters for various microbial species. Empirical cumulative distribution functions (*ECDFs*) for paired differences of between-(btwn) and within-pool (w/n) mutational and recombinational divergences (*θ*_*pool*_ and *ϕ*_*pool*_, respectively) and relative recombination rate of the pool ((*γ*/*μ*)_*pool*_). For each point, [*θ*_*pool*_(*btwn*) − *θ*_*pool*_(*w*/*n*)] = *θ*_*pool*_(*i*, *j*) − *θ*_*pool*_(*i*), where *θ*_*pool*_(*i*, *j*) is the between-pool divergence inferred for the between-cluster profile of clusters *i* and *j*, and *θ*_*pool*_(*i*) is the within-pool divergence for the within-cluster profile for cluster *i*. If either *θ*_*pool*_(*i*) or *θ*_*pool*_(*j*) could not be inferred, the paired difference was excluded. The same is true for the *ϕ*_*pool*_ and (*γ*/*μ*)_*pool*_ distributions. Additionally, only species where at least half of the clusters are represented in the *ECDF* are shown (i.e., recombination parameters could be inferred within-pool for at least half of the clusters in the dataset). Red circles indicate medians.

**Supplementary Fig. 4.**
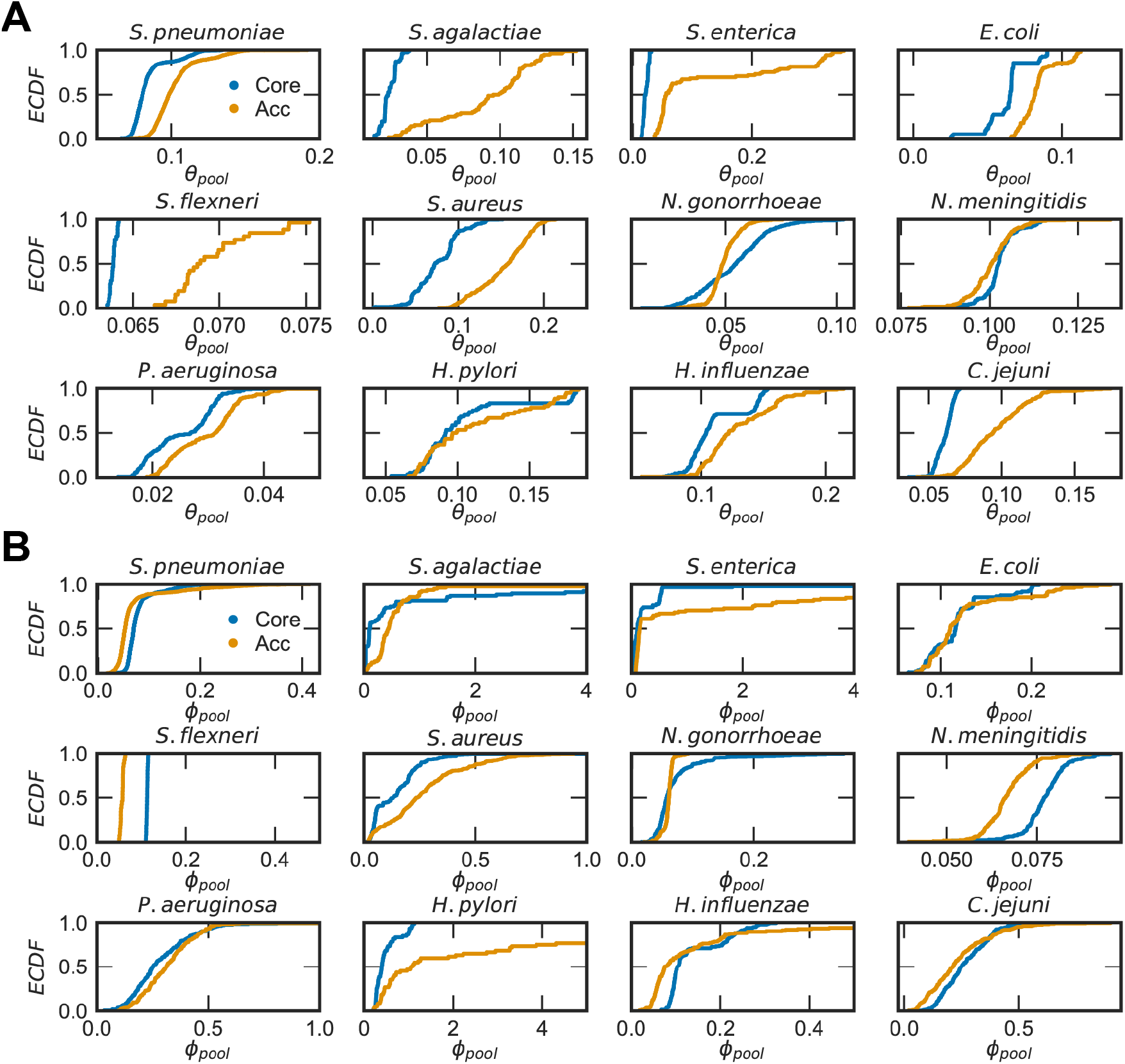
Empirical cumulative distribution functions (*ECDFs*) for the mutational and recombinational divergence for the 12 species shown in Fig. 3. (A) mutational divergence (*θ*_*pool*_), (B) recombinational divergence (*ϕ*_*pool*_). Both quantities are unitless. Core = core genes, Acc = accessory genes.

**Supplementary Fig. 5.**
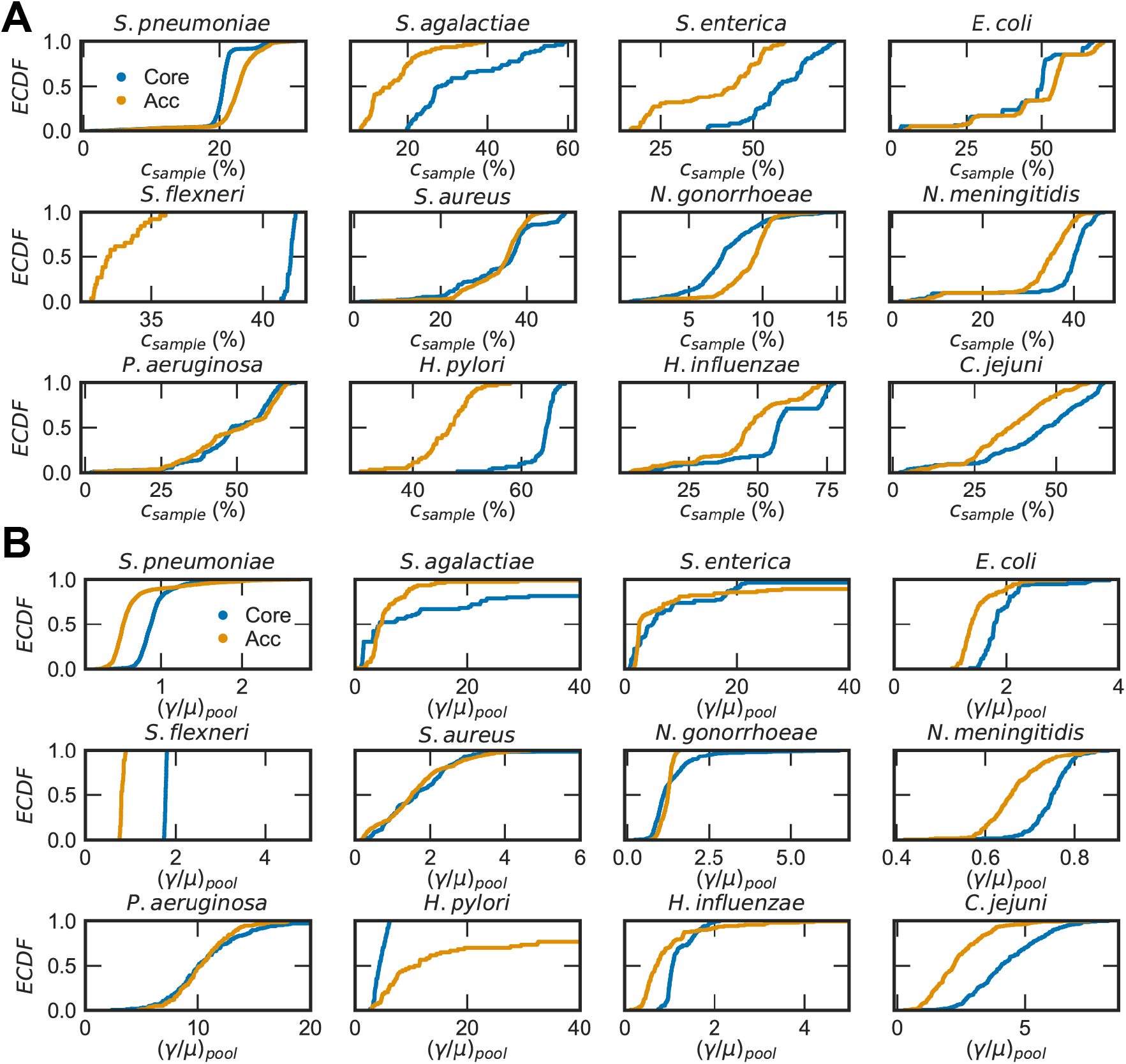
Empirical cumulative distribution functions (*ECDFs*) for the relative recombination rate and recombinational coverage for the 12 species shown in Fig. 3. (A) Relative recombination rate, (*γ*/*μ*)_*pool*_, which is unitless. (B) Recombinational coverage (*c*_*sample*_) given as the percentage of the genome which has been affected by recombination. Core = core genes, Acc = accessory genes.

**Supplementary Fig. 6.**
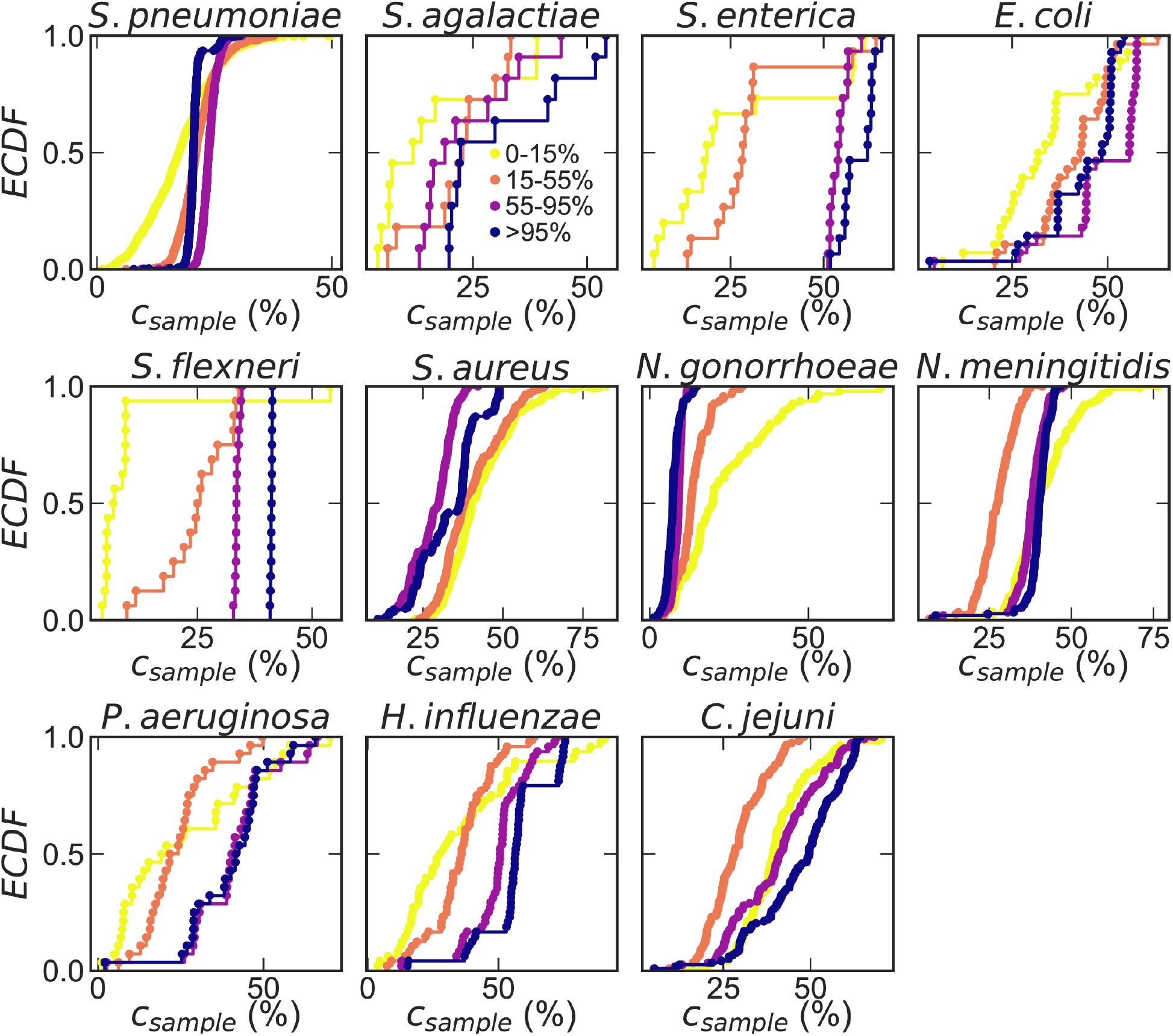
Empirical cumulative distribution functions (*ECDFs*) of recombinational coverage for gene frequency bins. Matched distributions of recombinational coverage (*c*_*sample*_, given as the percentage of recombined genomic sites) for gene frequency bins corresponding to Fig. 4. For each frequency range, upper bounds are inclusive, lower bounds exclusive.

**Supplementary Table 1.** Results of Wilcoxon signed-rank test comparing paired distributions for each of the 12 microbial species. Two-sided p-values were calculated using the Wilcoxon signed-rank test to compare the paired distributions of the mutational divergence (*θ*_*pool*_), normalized recombination rate ((*γ*/*μ*)_*pool*_), and recombinational coverage (*c*_*sample*_) for the core and accessory genome within each microbial species shown in Fig. 3A-C. Null hypothesis is that the distribution of paired differences is symmetric about zero (where *X*^*i*^(*acc*) − *X*^*i*^(*core*) is the paired difference for recombination parameter *X* inferred for the core and accessory genome of cluster or cluster pair *i*).

| Species | p-value ($\theta_{pool}$) | p-value ($(\gamma/\mu)_{pool}$) | p-value ($c_{sample}$) |
| --- | --- | --- | --- |
| *S. pneumoniae* | 1.92E-159 | 3.49E-99 | 5.02E-143 |
| *S. agalactiae* | 5.81E-14 | 2.93E-02 | 2.28E-13 |
| *S. enterica* | 1.17E-15 | 5.04E-01 | 1.14E-14 |
| *E. coli* | 2.60E-17 | 2.34E-16 | 2.57E-16 |
| *S. flexneri* | 8.30E-06 | 8.30E-06 | 8.30E-06 |
| *S. aureus* | 3.97E-65 | 2.61E-02 | 6.59E-01 |
| *N. gonorrhoeae* | 8.63E-06 | 6.98E-01 | 2.40E-31 |
| *N. meningitidis* | 3.04E-05 | 2.00E-23 | 2.17E-26 |
| *P. aeruginosa* | 9.24E-26 | 9.18E-01 | 7.29E-01 |
| *H. pylori* | 9.76E-02 | 2.84E-11 | 1.63E-11 |
| *H. influenzae* | 8.40E-20 | 1.32E-11 | 7.18E-24 |
| *C. jejuni* | 3.44E-36 | 1.70E-29 | 5.77E-31 |

**Supplementary Table 2.** Description of the microbial species analyzed in Figs. 2-4. Column descriptions are as follows: *“*species” and “full name” give the abbreviated species name used in the main text and full species name respectively, “major clusters” gives the number of sequence clusters analyzed, “total strains aligned” gives the number of consensus genomes (or whole genome sequences) which were made by aligning raw reads to the reference genome, “total strains in major clusters” gives the number of strains included in the major clusters (and therefore included in the analysis of recombination parameters), and “min strains per cluster” gives the minimum number of strains a cluster had to have to be designated as a major cluster. The minimum cluster size was lowered for smaller strain collections as follows (where *s* is the number of strains in the collection): for *s* > 10,000 in the collection the minimum cluster size was 100, for 10,000 > *s* > 5,000 the minimum was 50 (*S. enterica* was on the border, so we took the minimum to be 50), for 5,000 > *s* > 1,000 the minimum was 25, and for *s* < 1,000 the minimum was 10. The last four columns give the average synonymous diversity (*d*_*s*_) within clusters and between clusters (shown as mean +/− st. dev.).

| Species | Full name | major clusters | total strains aligned | total strains in major clusters | min strains per cluster | $d_s$ for core within cluster | $d_s$ For core between clusters | $d_s$ for flex within cluster | $d_s$ for flex between clusters |
| --- | --- | --- | --- | --- | --- | --- | --- | --- | --- |
| *S. pneumoniae* | *Streptococcus pneumoniae* | 44 | 26,599 | 24,097 | 100 | $(6.7 \pm 3.0) \times 10^{-3}$ | $(1.7 \pm 0.3) \times 10^{-2}$ | $(8.4 \pm 3.2) \times 10^{-3}$ | $(2.1 \pm 0.4) \times 10^{-2}$ |
| *S. agalactiae* | *Streptococcus agalactiae* | 17 | 7,917 | 7,365 | 50 | $(1.2 \pm 0.9) \times 10^{-4}$ | $(6.7 \pm 3.0) \times 10^{-3}$ | $(1.0 \pm 1.0) \times 10^{-3}$ | $(1.0 \pm 0.3) \times 10^{-2}$ |
| *S. enterica* | *Salmonella enterica* subsp. *enterica* | 18 | 10,357 | 6,296 | 50 | $(4.2 \pm 2.6) \times 10^{-5}$ | $(1.2 \pm 0.4) \times 10^{-2}$ | $(1.8 \pm 1.6) \times 10^{-4}$ | $(2.7 \pm 1.1) \times 10^{-2}$ |
| *E. coli* | *Escherichia coli* | 15 | 8,564 | 5,362 | 50 | $(4.9 \pm 2.0) \times 10^{-5}$ | $(2.6 \pm 1.5) \times 10^{-2}$ | $(7.9 \pm 0.1) \times 10^{-4}$ | $(3.5 \pm 1.8) \times 10^{-2}$ |
| *S. flexneri* | *Shigella flexneri* | 15 | 2,248 | 1,556 | 25 | $(2.0 \pm 0.8) \times 10^{-5}$ | $(6.8 \pm 0.1) \times 10^{-3}$ | $(8.7 \pm 7.8) \times 10^{-5}$ | $(6.6 \pm 9.0) \times 10^{-3}$ |
| *S. aureus* | *Staphylococcus aureus* | 30 | 25,125 | 22,535 | 100 | $(1.0 \pm 0.6) \times 10^{-5}$ | $(2.4 \pm 0.1) \times 10^{-2}$ | $(8.2 \pm 4.0) \times 10^{-3}$ | $(4.3 \pm 1.3) \times 10^{-2}$ |
| *N. gonorrhoeae* | *Neisseria gonorrhoeae* | 25 | 7,818 | 6,724 | 50 | $(6.6 \pm 4.2) \times 10^{-4}$ | $(3.9 \pm 0.7) \times 10^{-3}$ | $(1.0 \pm 0.5) \times 10^{-3}$ | $(4.6 \pm 0.6) \times 10^{-3}$ |
| *N. meningitidis* | *Neisseria meningitidis* | 18 | 13,017 | 10,968 | 100 | $(5.8 \pm 2.3) \times 10^{-3}$ | $(3.9 \pm 0.4) \times 10^{-2}$ | $(7.1 \pm 2.6) \times 10^{-3}$ | $(3.2 \pm 0.3) \times 10^{-2}$ |
| *P. aeruginosa* | *Pseudomonas aeruginosa* | 18 | 873 | 667 | 10 | $(1.9 \pm 2.2) \times 10^{-3}$ | $(1.3 \pm 0.4) \times 10^{-2}$ | $(2.3 \pm 2.5) \times 10^{-3}$ | $(1.5 \pm 0.5) \times 10^{-2}$ |
| *H. pylori* | *Helicobacter pylori* | 11 | 401 | 298 | 10 | $(3.9 \pm 0.3) \times 10^{-2}$ | $(6.2 \pm 1.7) \times 10^{-2}$ | $(2.8 \pm 3.5) \times 10^{-3}$ | $(4.7 \pm 1.2) \times 10^{-2}$ |
| *H. influenzae* | *Haemophilus influenzae* | 10 | 636 | 555 | 10 | $(1.4 \pm 0.8) \times 10^{-2}$ | $(6.3 \pm 2.1) \times 10^{-2}$ | $(1.1 \pm 0.6) \times 10^{-2}$ | $(5.9 \pm 2.1) \times 10^{-2}$ |
| *C. jejuni* | *Campylobacter jejuni* | 20 | 15,702 | 13,793 | 100 | $(2.4 \pm 0.3) \times 10^{-3}$ | $(2.8 \pm 0.8) \times 10^{-2}$ | $(8.0 \pm 3.6) \times 10^{-3}$ | $(3.2 \pm 0.7) \times 10^{-2}$ |

**Supplementary Table 3.** Results of Friedman test comparing matched distributions for each of the nine microbial species shown in Fig. 4. Two-sided p-values were calculated using the Friedman test to compare the matched distributions of the normalized recombination rate ((*γ*/*μ*)_*pool*_) and recombinational coverage (*c*_*sample*_) for all four gene frequency bins within each microbial species shown in Fig. 4. Null hypothesis in this case is that the measurements have been drawn from the same distribution.

| Species | p-value $((\gamma/\mu)_{pool})$ | p-value $(c_{sample})$ |
| --- | --- | --- |
| *S. pneumoniae* | 4.11E-97 | 1.76E-173 |
| *S. agalactiae* | 3.23E-02 | 5.31E-03 |
| *S. enterica* | 1.14E-01 | 4.15E-04 |
| *E. coli* | 2.09E-03 | 5.04E-08 |
| *S. flexneri* | 5.79E-09 | 1.00E-08 |
| *S. aureus* | 9.32E-03 | 2.44E-85 |
| *N. gonorrhoeae* | 4.53E-08 | 6.57E-36 |
| *N. meningitidis* | 5.41E-67 | 2.03E-63 |
| *P. aeruginosa* | 7.81E-02 | 1.00E-07 |
| *H. influenzae* | 1.82E-06 | 2.08E-19 |
| *C. jejuni* | 1.51E-18 | 1.48E-37 |

**Supplementary Table 4.** Identifying information for sequencing data used in study. Source of data and corresponding identifiers for all sequencing data used in this study.

| Type of data | Data source | Identifier(s) |
| --- | --- | --- |
| PubMLST database for microbial species analyzed | PubMLST | https://pubmlst.org/ |
| Reference genome for *Streptococcus pneumoniae* | NCBI RefSeq | RefSeq ID:<br>NC_003098.1;<br>https://www.ncbi.nlm.nih.gov/genome/176?genome_assembly_id=299864 |
| Reference genome for *Staphylococcus aureus* | NCBI RefSeq | RefSeq ID:<br>NC_007795.1;<br>https://www.ncbi.nlm.nih.gov/genome/154?genome_assembly_id=299272 |
| Reference genome for *Streptococcus agalactiae* | NCBI RefSeq | RefSeq ID:<br>NZ_LR134512.1;<br>https://www.ncbi.nlm.nih.gov/genome/?term=Streptococcus+agalactiae |
| Reference genome for *Salmonella enterica* subsp. *Enterica* | NCBI RefSeq | RefSeq ID:<br>NC_003197.2;<br>https://www.ncbi.nlm.nih.gov/genome/152?genome_assembly_id=305903 |
| Reference genome for *Escherichia coli* | NCBI RefSeq | RefSeq ID:<br>NC_000913.3;<br>https://www.ncbi.nlm.nih.gov/genome/?term=escherichia+coli |
| Reference genome for *Shigella flexneri* | NCBI RefSeq | RefSeq ID:<br>NC_004337.2;<br>https://www.ncbi.nlm.nih.gov/genome/?term=shigella+flexneri%5Borgn%5D |
| Reference genome for *Neisseria gonorrhoeae* | NCBI RefSeq | RefSeq ID:<br>NZ_CP012028.1;<br>https://www.ncbi.nlm.nih.gov/genome/864?genome_assembly_id=233772 |
| Reference genome for *Neisseria meningitidis* | NCBI RefSeq | RefSeq ID:<br>[NZ_CP021520.1;](https://www.ncbi.nlm.nih.gov/genome/172)<br>https://www.ncbi.nlm.nih.gov/genome/172 |
| Reference genome for *Pseudomonas aeruginosa* | NCBI RefSeq | RefSeq ID:<br>[NC_002516.2;](https://www.ncbi.nlm.nih.gov/genome/?term=txid287[orgn])<br>https://www.ncbi.nlm.nih.gov/genome/?term=txid287[orgn] |
| Reference genome for *Haemophilus influenzae* | NCBI RefSeq | RefSeq ID:<br>[NZ_CP009610.1;](https://www.ncbi.nlm.nih.gov/genome/?term=haemophilus+influenzae%5Borgn%5D)<br>https://www.ncbi.nlm.nih.gov/genome/?term=haemophilus+influenzae%5Borgn%5D |
| Reference genome for *Campylobacter jejuni* | NCBI RefSeq | RefSeq ID:<br>[NC_002163.1;](https://www.ncbi.nlm.nih.gov/genome/?term=Campylobacter+jejuni)<br>https://www.ncbi.nlm.nih.gov/genome/?term=Campylobacter+jejuni |
| XMFA for *Helicobacter pylori* | Ref 28 from main text; Dryad data repository | DOI:<br>[10.5061/dryad.8qp4n](https://doi.org/10.5061/dryad.8qp4n) |
| Raw reads corresponding to whole genome sequences for *Escherichia coli* and *Shigella flexneri* | NCBI, Bioproject: "Routine surveillance of E. coli and Shigella by Public Health England" | Bioproject ID:<br>PRJNA315192;<br>https://www.ncbi.nlm.nih.gov/bioproject/?term=PRJNA315192 |
| Raw reads corresponding to whole genome sequences for *Haemophilus influenzae* | Ref <sup>63</sup>; NCBI, Bioproject: "WGS Data for Haemophilus influenzae isolates" | Bioproject ID:<br>PRJNA512636;<br>https://www.ncbi.nlm.nih.gov/bioproject/?term=PRJNA512636 |

